# The role of long-range transcriptional regulation in interpretation of non-coding variants associated with human disease

**DOI:** 10.64898/2026.06.15.731051

**Authors:** Katarina Mandić, Dalibor Hršak, Filip Uljanić, Boris Lenhard, Anja Barešić

**Affiliations:** Division of Computing and Data Science, Ruđer Bošković Institute, Bijenička cesta 54, Zagreb, 10,000, Croatia; Institute of Clinical Sciences, Faculty of Medicine, Imperial College London, London, W12 0HS, United Kingdom; Laboratory of Medical Sciences (LMS), Medical Research Council (MRC), London, W12 0HS, United Kingdom

**Keywords:** genomic regulatory blocks, variant target gene, long-range transcription regulation, non-coding variants

## Abstract

Genome-wide association studies (GWAS) are the key tools for the discovery of associations between single nucleotide polymorphisms (SNPs) and phenotypic traits and have been successfully applied to many diseases and disorders. However, a great challenge is to find the gene affected by the non-coding fraction of SNPs, especially if the gene is distal in terms of genomic distance. In this study, we present a novel approach, named targPred, which utilises genomic regulatory blocks (GRBs) for inference of a connection between a certain SNP/locus and the target gene located in the same GRB, in a more robust and generalisable manner. We identified that many disease traits such as cancer and psychiatric disease have a propensity for long-range regulation. Furthermore, we showcased a childhood obesity locus which is connected to the distal *BDNF* gene. Finally, we propose a new web-based service based on enhancer-promoter association, to facilitate finding the causal genes for a wide array of traits and conditions.

## 1 Introduction

The complexity of molecular functions in an organism is maintained through dynamic mechanisms of gene expression activity regulation, as well as the conservation of those sets of genomic variants that have proven to optimize the organism’s genetic fitness. Transcriptional activity of individual genes is regulated by non-coding elements, sometimes in genomic vicinity of the gene promoter, but often across long distances in the genome’s primary sequence, ranging up to 1 megabase (Mb) or more [1– 3]. This type of regulation is termed long-range regulation (LRR) and is achieved through three-dimensional DNA folding and chromatin looping, bringing the non-coding elements into close physical proximity of the gene promoter, and regulating gene expression through intermediary proteins and transcription factors [4, 5]. Chromatin capture methods identify chromatin spatial arrangements by defining genomic regions with most mutual physical contacts such as topologically associating domains (TADs) [6, 7]. The disruptions in TADs generate novel and unpredictable enhancer-promoter interactions and erroneous gene activation/silencing events [8, 9], which can cause a myriad of diseases and disorders of neurological, psychiatric, developmental or oncogenic nature [10, 11]. All regulatory elements are hereafter termed enhancers, irrespective of their enhancing or repressive direction of effect on the transcriptional activity of the gene.

Within the genomes of higher order organisms, regions with a high density of evolutionarily conserved non-coding segments were identified, separately evolving in the invertebrate and vertebrate lineage [12–14]. These non-coding segments are found to be exceptionally well conserved through the phylogenetic system in terms of spatial arrangements, gene order (synteny), enhancer-promoter relationships (i.e. promoters maintain their associated sets of enhancers) and other functional characteristics [15, 16]. These regions, named genomic regulatory blocks (GRBs), house key developmental genes and complex regulatory landscapes which may span several megabases [17]. The GRB provides a model for the association of enhancer-promoter pairs which operate through LRR across large genomic distances ensuring a precise control of developmentally important target genes, while bystander genes are, by and large, unresponsive to these elements [17, 18]. Moreover, GRBs generally correspond well with individual TADs, which indicates a shared underlying mechanisms of evolutionary conservation for both phenomena [19].

Finding the causal links between the SNPs/loci and health outcomes is a target of active research, as it leads to discovery of molecular mechanisms targetable by various therapeutics. Genome-wide association studies (GWAS) are the key tools for the discovery of associations between SNPs and phenotypic traits and have been successfully applied to many diseases and disorders [20–22]. More than 90 % of GWAS SNPs are non-coding and they are often located in regulatory regions distant from a coding gene [23–25]. The clinically prevalent approach of assigning the nearest gene as a target fails to provide mechanistic insights into disease aetiology and the gene’s functional relevance, as well as pleiotropic and epistatic effects.

To address this issue, several methods were recently developed, combining experimental data on chromatin contacts with various genomic function predictions. These integrative SNP-to-gene (S2G) frameworks combine 3D contacts with functional chromatin maps such as DNase-seq or ATAC-seq accessibility and histone modifications to infer enhancer–promoter interactions and link variants to genes. A prominent example is the Activity-by-Contact (ABC) model, which defines an element’s contribution to activity of a gene as the product of its biochemical “activity” (quantified from DNase/ATAC and H3K27ac ChIP-seq signal) and its Hi-C contact frequency with the gene’s promoter, and was trained and validated using CRISPRi-FlowFISH perturbation screens to predict enhancer–gene connections genome-wide [26]. In parallel, the Open Targets Genetics locus-to-gene (L2G) model employs gradient-boosted machine learning to integrate fine-mapped GWAS credible sets with distance, molecular QTL colocalisation, chromatin interaction and variant pathogenicity features, yielding an L2G score that ranks likely causal genes at each locus within the Open Targets Platform [27]. Both ABC and L2G are trained on curated sets of “gold-standard” loci and phenotypes which, although carefully assembled, remain relatively limited in size and biased toward well-studied traits, potentially constraining the generalisability of these models across tissues and disease areas. Complementary resources such as GeneHancer integrate multiple evidence streams (including enhancer RNA co-expression, eQTLs and promoter-capture Hi-C) to assign likelihood-based scores to enhancer–gene pairs and define “elite” regulatory interactions with stronger support [28]. Beyond individual resources, a recent heritability-based framework systematically evaluated and linearly combined diverse S2G strategies, showing that an optimal combined model (cS2G) incorporating exon, promoter, fine-mapped cis-eQTL, enhancer-gene linking (EpiMap, ABC) and chromatin accessibility-derived strategies more than doubles recall at fixed high precision, outperforming previous integrated approaches such as GeneHancer and Open Targets in terms of disease heritability explained [29]. Nevertheless, most existing frameworks remain fundamentally locus-centric and are trained within a single-species regulatory landscape, often underrepresenting long-range architectural constraints and evolutionary conservation of regulatory wiring. Motivated by these gaps, we develop a GRB-based approach, that explicitly leverages evolutionarily stable GRBs and focuses on long-range interactions, to associate non-coding variants with genes across diverse traits in a more robust and generalisable manner.

Here we present targPred, a method that enables inference of a connection between a certain SNP/locus and a phenotypic trait by observing the association between the regulatory enhancers within a locus and the target gene located in the same GRB. These enhancer-promoter associations can be readily observed from tissue-wide transcriptional activity profiles, available in FANTOM5 [30, 31]. The GRB-based approach bears the advantage over methods relying on chromatin contacts and functional maps in the reduced costs of obtaining data, as identifying GRBs requires data on evolutionary preservation of a genomic region, in turn providing more tissue and cell-type invariant information on regulatory patterns. This approach has previously been successfully applied to disorders such as type II diabetes and obesity [32], and has been used to identify genes under LRR responsible for a set of neuropsychiatric disorders [33].

In this paper, we extend the scope of the methodology outlined in the work by Barešić et al. from neuropsychiatric disorders to all phenotypic traits listed in the continually updated GWAS catalogue [25]. We also present a new web-based service named targPred (http://targpred.irb.hr/) intended to facilitate finding the causal genes to a wide array of traits and conditions by using targPred. The main advantage we see is that our method does not rely on a large number of datasets, feature extraction or the inclusion of epistatic effects, as well as providing the capability of detecting the LRR effects, not systematically attainable by other methods. We aim to make our service a valuable resource for both the scientific community and future clinical practices, as well as creating a base for expansion of the knowledge on the long range gene regulation.

## 2 Results

We applied targPred for assessing candidate genes for 280,711 variants, across 11,921 phenotypic traits from the GWAS catalog yielding 757,253 autosomal and X-chromosomal GWAS SNP-trait associations, further supporting these associations with a statistical framework for enhancer-gene links. Hereafter, we define a GWAS locus as a region around the tagging SNP which is in linkage disequilibrium (LD). targPred resource reports the GRB target gene as a plausible alternative target to the GWAS-proposed genes for the 38 % of loci located within a GRB (Figure 1A, middle ring). Finally targPred provides novel gene candidates with respect to GWAS reported and mapped genes for 158,611 locus-trait pairs comprising 56% of all pairs located in the GRBs (Figure 1A, shown in yellow).

**Fig. 1.**
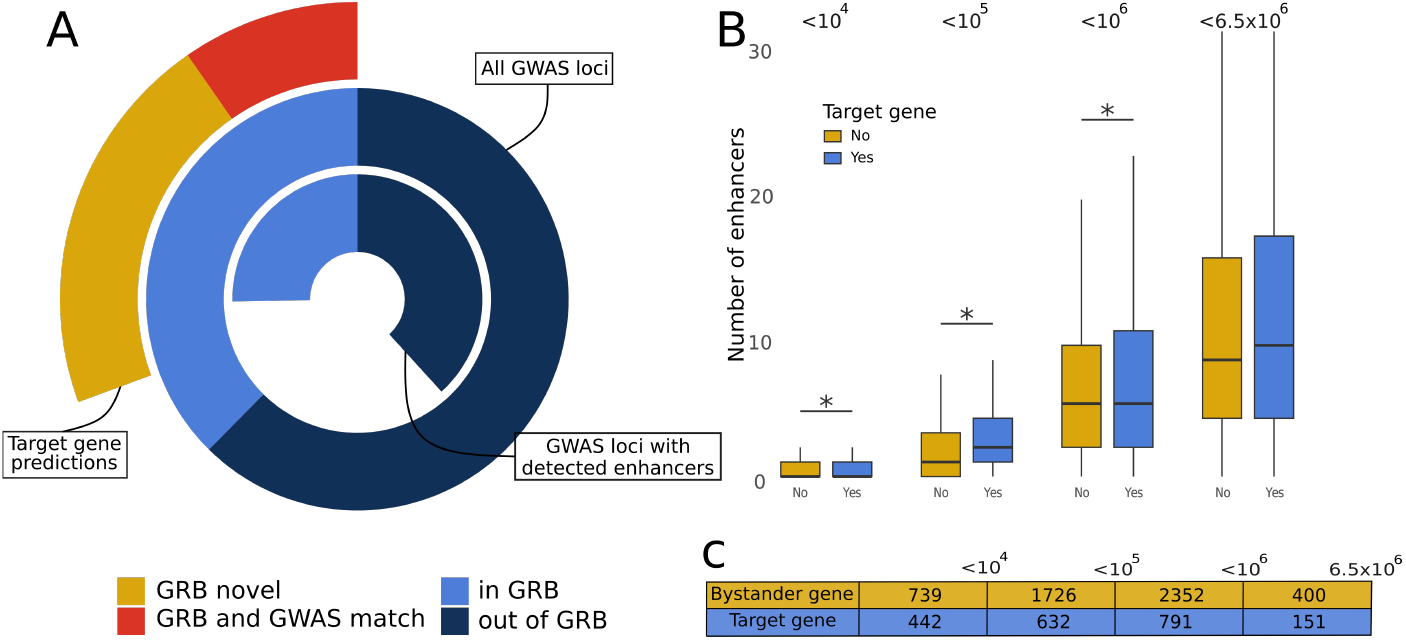
**A** Middle ring shows all GWAS loci. The inner ring shows loci which overlap enhancers. The outer ring shows SNP-gene assignments based on targPred - red are loci for which targPred assigns the same gene as the GWAS catalog, yellow are loci for which targPred assigns novel target genes. **B** Cumulative count of enhancers significantly associated (BH adjusted empP < 0.05) with bystander and target genes binned by genomic distance. **C** Number of bystander and target genes with significant associations per bin.

### 2.1 GRB target genes are more responsive to LRR

Additionally, for a subset of GRB loci where a locus overlaps a known enhancer (shown in the inner ring in Figure 1A, 67 %), we provide an experimentally-grounded strength of association between eRNA activity and genes’ activity over a wide range of biosamples (cell lines, primary cells and tissues). We computed genome-wide enhancer-gene empirical p-values (empP) using the CAGE dataset published in FANTOM5 project [31]. These empP values are based on a permutation test of CAGE signals from enhancers and promoters across 309 biosamples, capturing both enhancing (increased expression) and repressive (decreased expression) scenarios (see method section for more detail). Significant empP values (Benjamini-Hochberg (BH), adjusted p < 0.05) indicate activity consistent with functional enhancer promoter interaction.

On average, GRB target genes have a higher count of locus-associated enhancers than bystander genes across different genomic distance bins (Figure 1B). This difference is significant in the *<* 10^4^, *<* 10^5^, and *<* 10^6^ bins (Wilcoxon rank-sum test, p<0.05) suggesting that target genes have a more complex regulatory landscape that spans a larger genomic range (Supplemental table 1).

### 2.2 Using the targPred resource

We provide targPred target gene reannotation in a publicly-available resource. The data obtained from GWAS, version 2026 [34] can be found in tab “All SNPs” containing all 757,253 SNP-trait associations. The information on the loci are located in the tab “All Loci” (174,376 different GWAS loci). The user can browse the tables by mapped trait, filter by whether associated loci are inside or outside a GRB, and search for specific overlapping genes. An example of the SNP view is given in Figure 2. Finally, the tab “Locus Summary” contains the results of the method described in the previous section: if a GWAS locus is within a GRB and overlaps one or more enhancers, a two-sided distribution plot matrix of log expression values (in TPMs) for each enhancer-gene pair and respective empirical p-values are shown, as in Figure 3. Therein, we present an example of a summary for the major depressive disorder and the GWAS locus chr5:141,208,305-141,210,753, which contains a SNP rs166040 [35]. This locus lies within the GRB region chr5:140,937,495-141,353,829, where a target gene named *RELL2* (Receptor Expressed in Lymphoid tissues-like 2, also denoted as *C5orf16* ) is also located, which displays a statistically significant increase in transcriptional activity in tissues where the eRNA signal is detected. To the best of our knowledge, this association is novel and this gene is relatively distant from the interacting enhancer chr5:141,209,314-141,209,613 (192,900 bp) located in the afore-mentioned locus. *RELL2* is a gene responsible for inducing cell apoptosis in tumour tissues by activating the MAPK14/p38 cascade and it is generally overexpressed in tumour tissues [36, 37]. The enrichment analysis has also revealed the participation of *RELL2* in a number of metabolic pathways associated with inflammation and potentially linked to depression, such as IL-6/JAK/STAT3 [38, 39] and Toll-like receptor pathway [40]. According to the GTEx portal, *RELL2* is most expressed in brain tissues (pituitary gland, cerebellum and cortex), spleen and testes [41]. Given the connection between inflammation and depression is widely reported and the above mentioned metabolic pathways are heavily involved in both processes [42–44], and the tissue specific expression profile of *RELL2*, our finding of a direct connection between the activity regulation of *RELL2* through the aforementioned enhancer and the occurrence of major depressive disorder has a solid foundation in the literature.

**Fig. 2.**
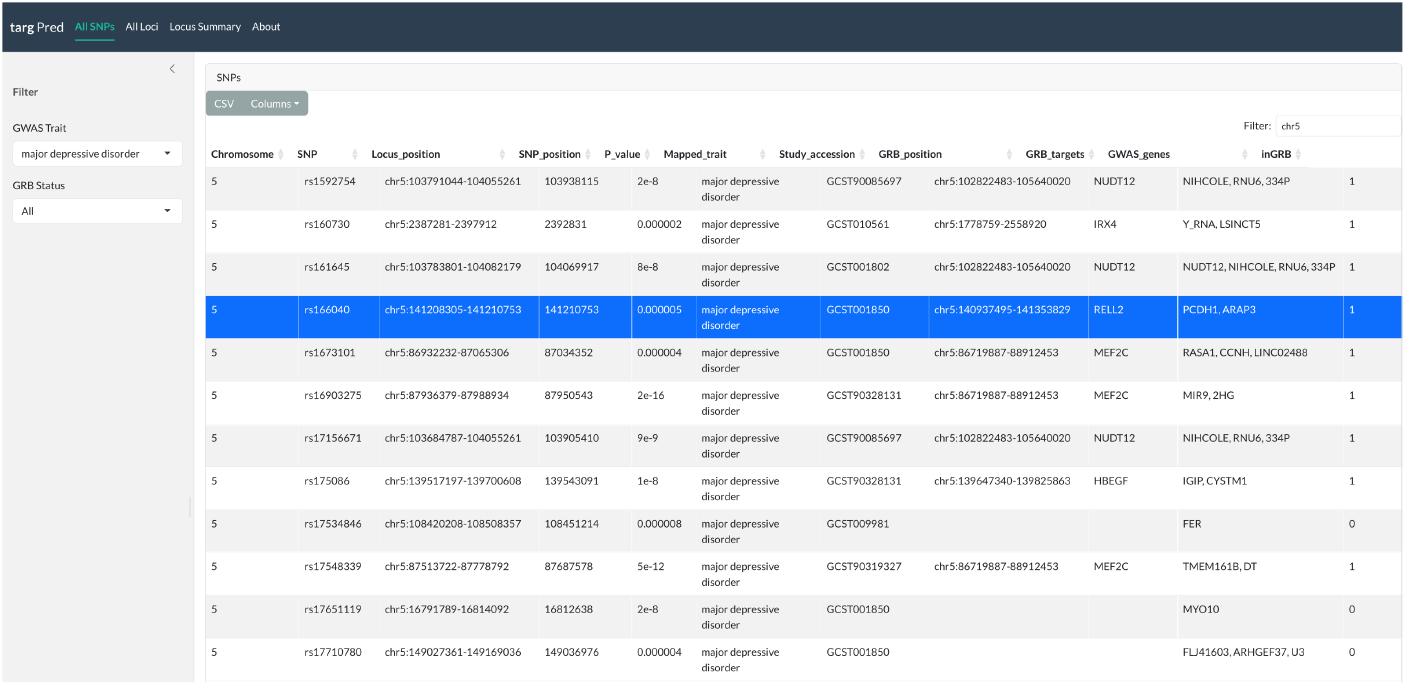
“All SNPs” tab of the targPred online resource, showing the interactive table with GRB data for SNPs from the GWAS Catalog.

**Fig. 3.**
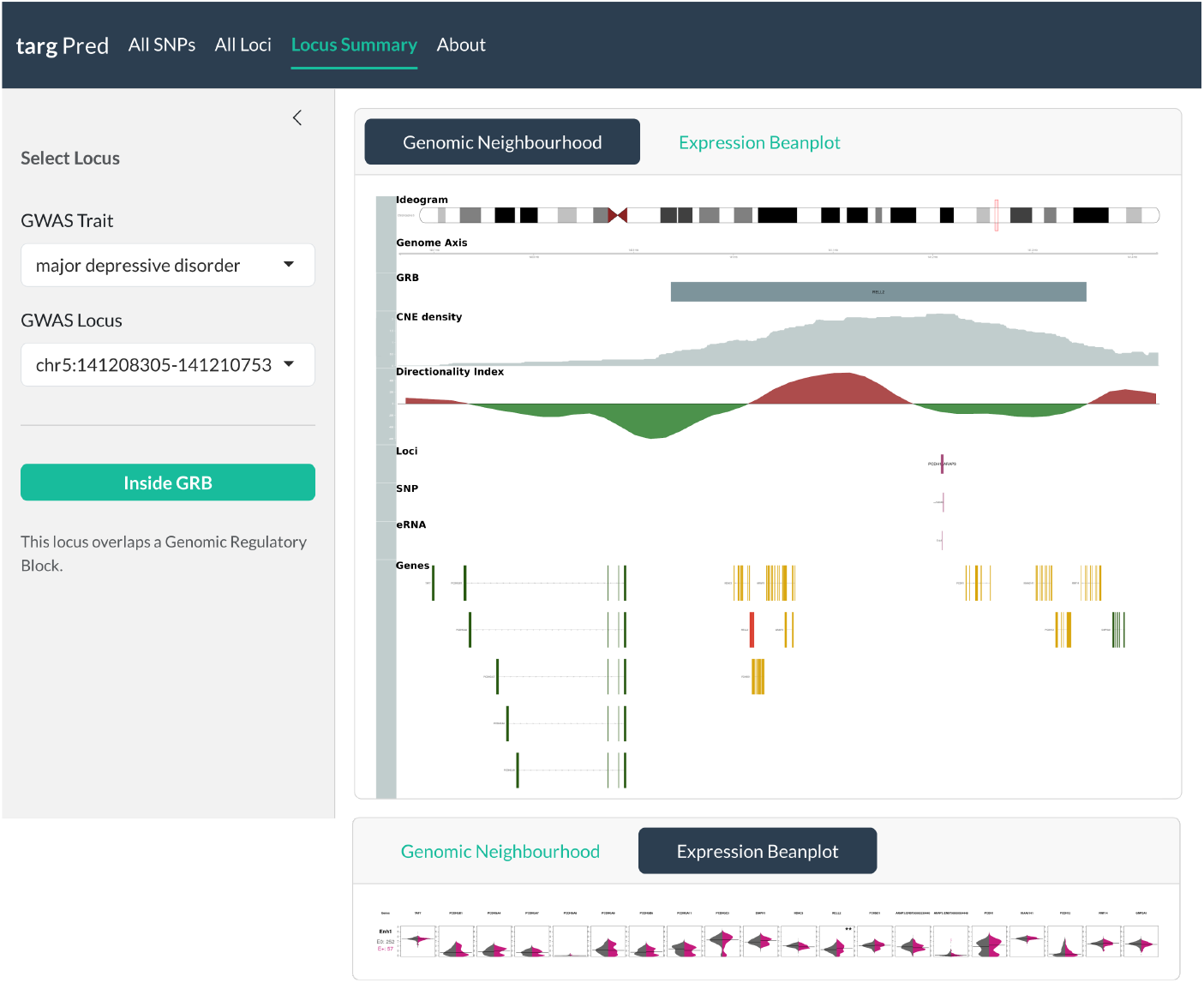
“Locus Summary” tab of the targPred online resource contains the genomic neighbourhood and statistical plots for enhancer-containing loci found within GRBs. The left side contains the general description of the locus as well as two drop-down menus for trait and loci selection. The main window consists of two options: Genomic Neighbourhood and Expression Beanplot. The Genomic Neighbourhood shows 9 genomic tracks for the GRB region where the locus is located; description for each track is found in Table 1. The Expression Beanplot contains a matrix of two-sided distribution plots depicting the association between eRNA and gene expression levels. For the sake of clarity, the plots have been rescaled and rearranged in this Figure.

### 2.3 Traits under LRR

We assessed LRR in traits represented in the GWAS, inspecting specifically loci that harbour enhancers. We compared trait-associated enhancer-gene pairs (empP < 0.001) that fall inside and outside GRB regions, and tested the genomic distance of these pairs using the Wilcoxon rank sum test. A total of 1607 disease traits have at least one locus in a GRB which contains an enhancer-gene pair. Among these, we found 779 traits that had significantly larger distances for enhancer-gene pairs inside GRBs versus outside GRBs (BH-adjusted p *<* 0.05). Top 30 disease traits sorted by p-value, further arranged by the median difference were selected to illustrate the results (Figure 4A). We observed six broad categories of disease traits: cancer (EFO term “cancer”, MONDO:0004992), immune (EFO term “immune system disorder”, MONDO:0005046), asthma (EFO term “asthma”, MONDO:0004979), metabolic (EFO term “metabolic disease”, MONDO:0005066), psychiatric (EFO term “psychiatric disorder”, MONDO:0002025) and other. Using this approach resulted in categorizing *Clostridium difficile* infection and oral lichen planus as “other”. However, these traits are connected to the immune and cancer categories since *Clostridium difficile* infection is strongly associated with inflammatory bowel disease [45] and oral lichen planus is a T-cell-mediated inflammatory disease strongly associating with autoimmune disease and oral cancer [46]. Additionally, among significant traits, we note a distinct group of psychiatric disorders found to be under LRR (Figure 4B), including a range of substance-related disorders, cognitive and mental disorders. Furthermore, we found enrichment of GWAS loci in GRBs using a genome permutation test, demonstrating that loci of some of the psychiatric disorders are more often observed in GRBs (Figure 4C). This was previously shown for schizophrenia and autism spectrum disorder [33]. Similar results were found in our study when using a different GWAS datasets for these traits. Additionally, we note enrichment of GWAS loci from a major depressive disorder study and a study combining 8 psychiatric disorders (Figure 4C).

**Fig. 4.**
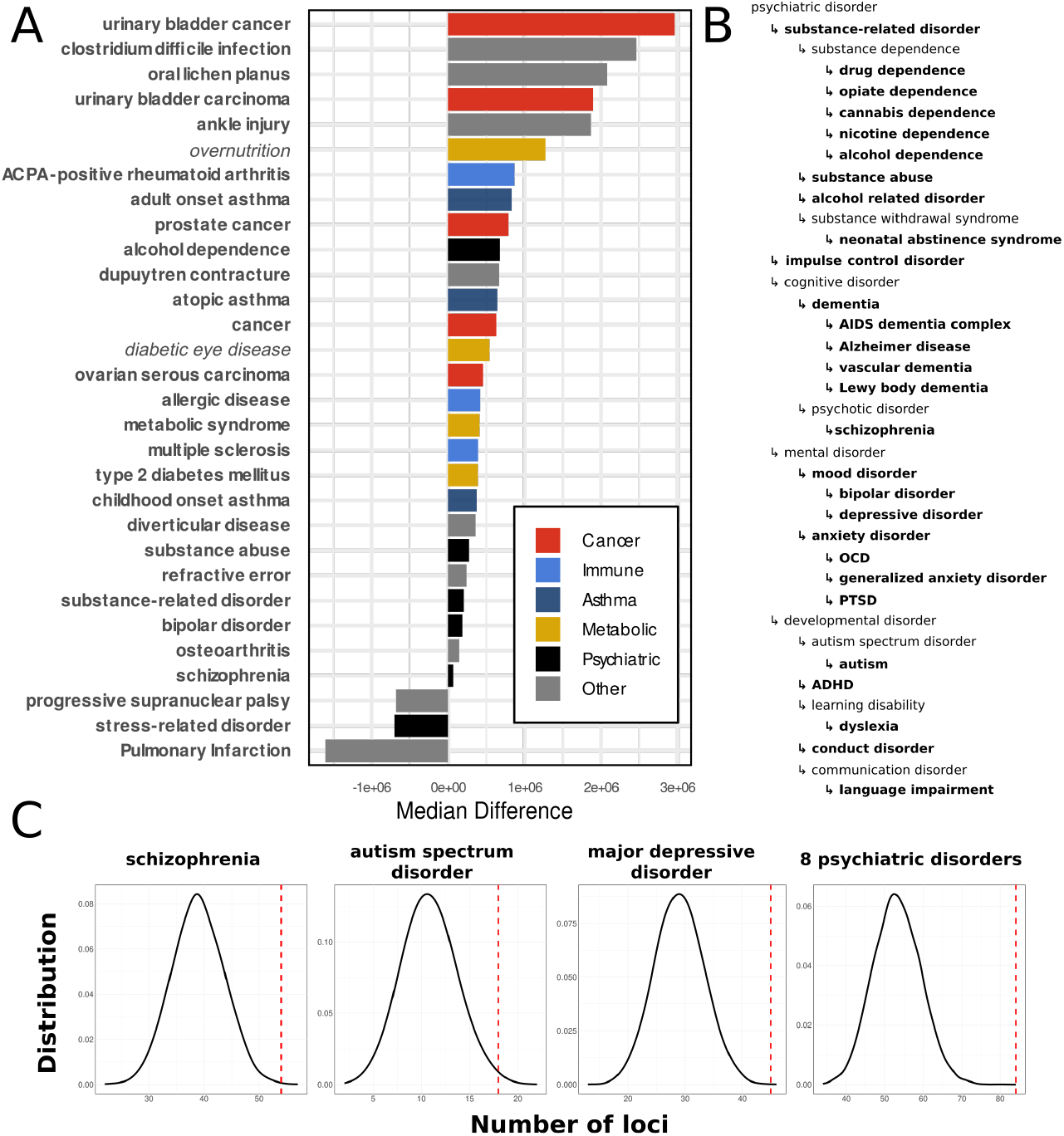
**A** Top 30 traits filtered by BH adjusted p-value with LRR in GRBs, and sorted by median distance of enhancer-gene pairs inside versus outside of GRBs. For the classification of italicised terms into metabolic class, see Methods section. **B** Overview of the psychiatric disorder ontology with high-lighted traits (bolded) which have significant LRR in GRB versus outside of GRB. Acronyms: OCD - obsessive-compulsive disorder; PTSD - Post-traumatic stress disorder; ADHD - attention deficit-hyperactivity disorder. **C** The probability distribution of loci overlapping N = 10,000 permutated GRB regions was estimated for four relevant psychiatric disorders. P-value were calculated empirically for the number of loci overlapping observed GRBs. All four disease traits show significant enrichment in GRBs (p-value < 0.05, see Supplemental table 2).

### 2.4 Comparing SNP-gene assignment methods

The targPred approach predicts target genes for disease-associated common variants and we used two approaches to validate these predictions based on ground-truth datasets, described below. First, a dataset of disease-associated coding variants from ClinVar was used as the ground-truth for S2G associations - for coding variants, these associations are considered precise and unambiguous. Second, a set of experimentally-validated S2G associations by Alsheikh et al. [47] was used. It is worth noting here that both sets are rather limited in amount of data and its representativeness of the entire phenotype complexity.

Finally, we validate the gene lists predicted by targPred against the lists produced by similar computational S2G predicting tools: GeneHancer, ABC and cS2G model [28, 29, 48]. All comparisons to other prediction methods were performed on the subset of enhancers falling within the GRB regions.

#### 2.4.1 Comparison with validated SNP-gene links

To validate targPred predictions against the ground-truth dataset, we used coding variants deposited in the ClinVar database. In our approach, we searched for cases where a gene had both a coding variant and a nearby (within a GRB) non-coding variant associated with the same disease trait by the ClinVar and the targPred respectively, resulting in 432 variants (Figure 5A, Supplemental table 3). targPred supports gene assignment for 201 variants, out of which 102 were not reported in the GWAS catalog. For example, two variants (rs55754224, rs6829664) associated with atrial fibrillation are localized in the intronic region of *CAMK2D* which is the assigned target gene in GWAS (Supplemental figure S1) [49–51]. However, targPred predicts ANK2 as the target gene which is associated with various cardiovascular phenotypes, including atrial fibrillation, long QT syndrome and cardiac arrhythmia, reported in the ClinVar database. This prediction indicates ANK2 as a promising potential target, highlighting its relevance for these non-coding variants, in the context of atrial fibrillation, as well as two additional cardiovascular traits.

**Fig. 5.**
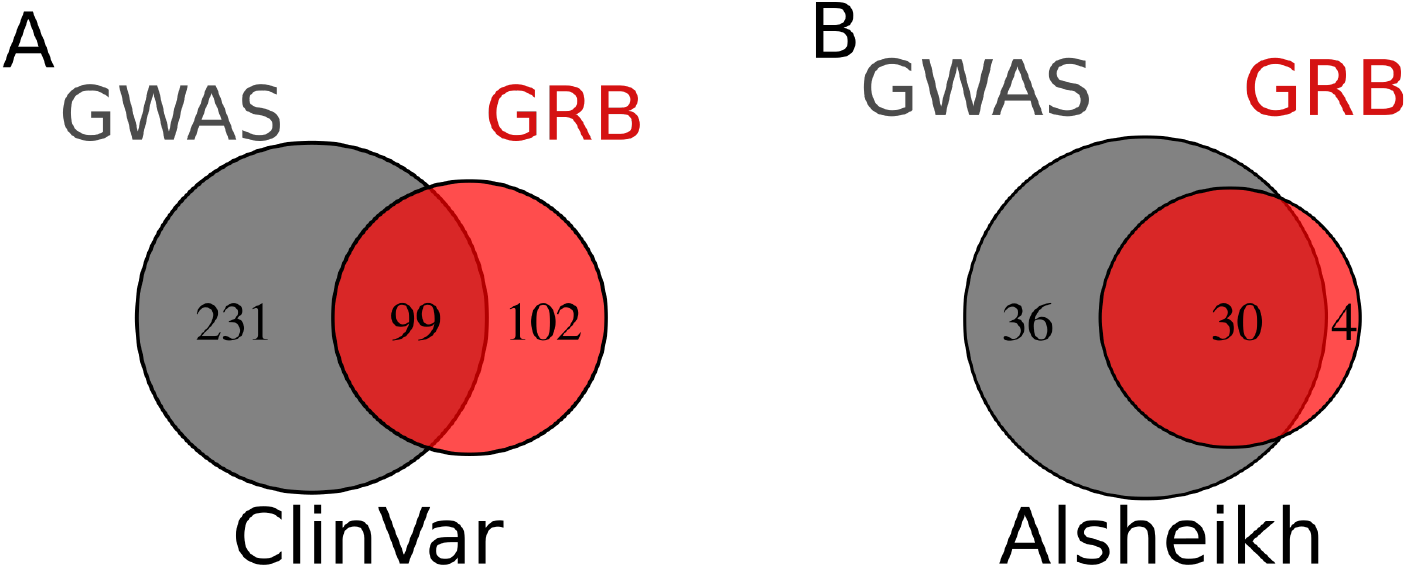
Comparison of GRB and GWAS target genes with experimentally-validated sets ClinVar [54] in **A** and Alsheikh [47] in **B**. The overlapping was marked as true if any of the predicted target genes on the given list were overlapping.

The second validation dataset originated from a list of manually curated publications providing experimental evidence for associations between regulatory variants and putative target genes [47]. On a subset of targPred associations presented by Alsheikh et al., the majority of SNP-gene links from the experimentally-validated dataset are supported with the same gene assignment in both the GRB and the GWAS target gene lists (Figure 5B, Supplemental table 3), with a small number of links unique to targPred. Therein, a variant (rs1859962) associated with prostate carcinoma assigned to *CASC17* a long non-coding RNA gene in GWAS [52, 53], however, targPred and the Alsheikh sets assign the variant to *SOX9*, which is over 1 Mb away.

#### 2.4.2 Comparison with other enhancer-gene predictive methods

Next, targPred enhancer-gene assignments were compared with ABC and GeneHancer enhancer-gene scores, and cS2G SNP-gene scoring. To this end, we defined three categories that each of the pairs reported in the 3 sets could fall into: 0 - the enhancer-gene pair is not significant in our dataset and does not include a GRB target gene, 1 - the enhancer-gene pair is significant in our dataset or includes a GRB target gene, 2 - the enhancer-gene pair is significant in our dataset and includes a GRB target gene. For comparison with the ABC set, we selected enhancer-gene pairs with scores > 0.015 as previously used [26] (Figure 6B) and for the GeneHancer set we selected pairs with scores > 250 due to a discontinuous distribution of the scores in the original dataset (Figure 6C). When comparing these representative predictive methods, we show that gene-enhancer pairs have statistically higher scores for category 2 in all the methods (Supplemental table 4). All three tested methods had higher scores in our category 2 than in 1 and 0 (significant increase for ABC and cS2G, not significant for GeneHancer, see Supplemental table 5). However, targPred identified many individual significant LRR enhancer-gene pairs that were missed by the methods listed above.

**Fig. 6.**
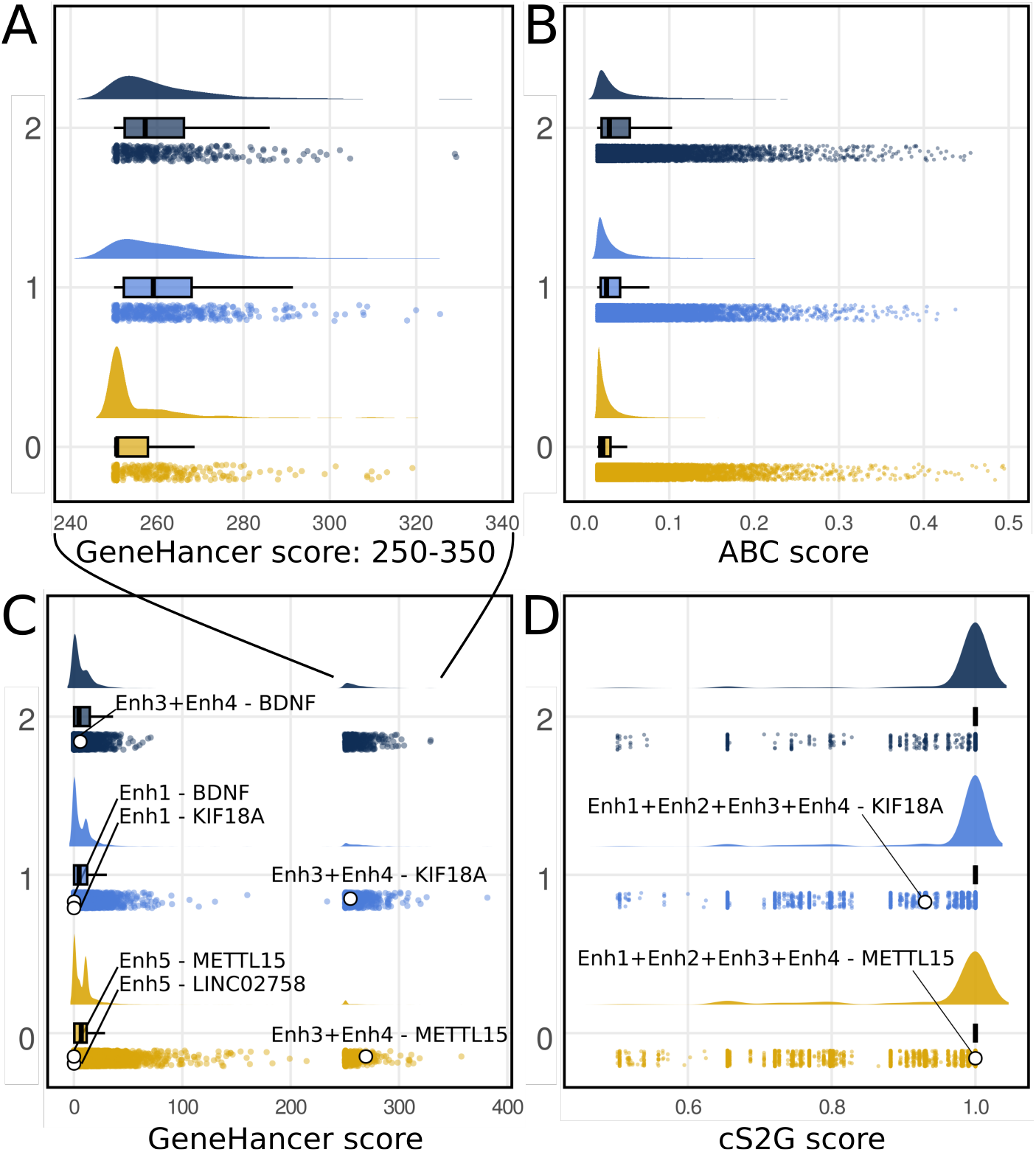
Comparison of GRB annotation with other S2G prediction methods: GeneHancer [28]) in **A** and **C**, ABC [48] in **B** and cS2G [29] in **D**. A distribution of GeneHancer scores is shown in **C** for the full range of scores, and **A** for the high-confidence scores >250. Individual scores for the enhancers from the childhood obesity loci are shown as white points in **C-D**. No ABC enhancers were found overlapping these two loci.

We next show a representative case for the *BDNF* target gene.

### 2.5 *BDNF* under long-range regulation in childhood obesity

Here we illustrate a locus-gene assignment using targPred, supported by the chromatin contact data for two childhood obesity loci. We have found that SNPs associated with obesity and other traits relating to the metabolic syndrome are enriched in GRBs (Supplemental figure S2, Supplemental table 2) and often form long range associations (Figure 4A). Figure 7A demonstrates two childhood obesity related loci (rs7940756 and rs10835310) overlapping a GRB (chr11:27,604,430-29,063,089). For these two loci the nearest genes are *METTL15* and *LINC02758* which were reported as targets in the original GWAS study [55]. However, targPred suggests a more plausible target gene *BDNF* supported by functional evidence, in contrast to the GWAS target gene prediction. *BDNF* was previously implicated in adult obesity, type 2 diabetes mellitus and other traits related to the metabolic syndrome. *METTL15* codes for a methyl-transferase protein, while *LINC02758* is a long non-coding RNA molecule with no clear function determined to date. The *METTL15* locus spans the entire *METTL15* and *KIF18A* genetic sequences, as well as four enhancers (Enh1-4, Figure 7A). Based on available transcriptional activity profiles, *BDNF* and *KIF18A* are responsive to the enhancer elements, while *METTL15* shows an expression profile of an ubiquitously expressed gene, largely unresponsive to the enhancer elements (Figure 7B). Chromatin contact map shows the region spanning the four enhancers interacts with the promoter region of *BDNF* effectively forming a regulatory hub [56]. The *LINC02758* locus spans one enhancer (Enh5) with no genes in the vicinity. targPred suggests that the *LINC02758* locus targets the distal *BDNF* gene across *∼* 1Mb distance. Furthermore, we found *BDNF* to be significantly responsive to Enh5 with Hi-C data showing that these two regions form chromatin contacts (shown with arrow, Figure 7A). This example also demonstrates how targPred may provide novel S2G associations, which are missed by recently published methods (Figure 6B-D).

**Fig. 7.**
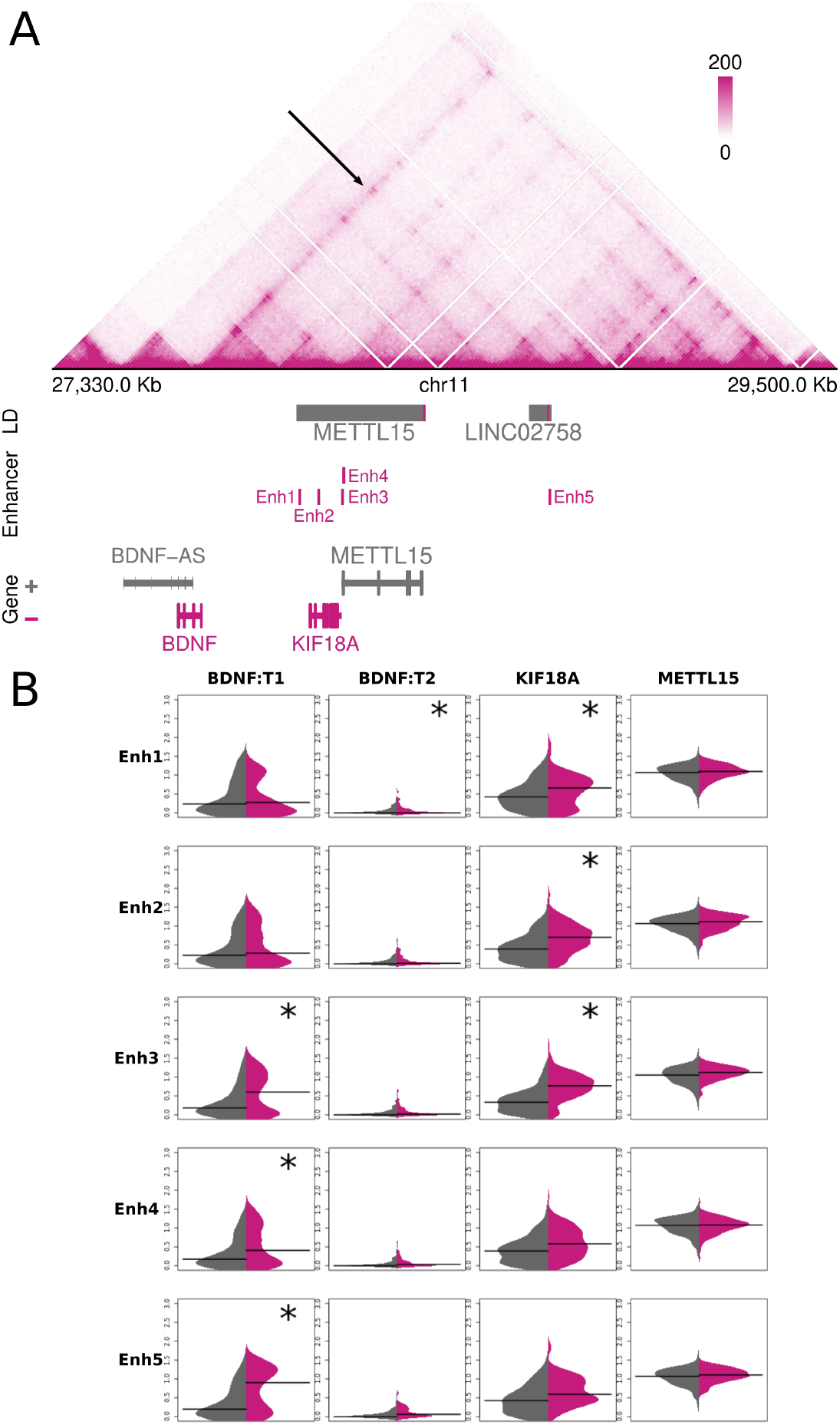
Childhood obesity loci associated with *BDNF*. **A** Hi-C map of the GRB which harbours the *BDNF* target gene with annotated loci relating to childhood obesity and an arrow highlighting a long-range interaction. Two loci are marked in grey, with accompanying SNPs shown overlapping in pink in the LD track and labeled with the GWAS target gene assignment for every locus. **B** Two-sided distribution plot matrix of log-TPM expression values for all enhancer–gene pairs in this GRB. The distribution of expression of each gene in tissues where the enhancer is inactive or active is shown in grey and pink, respectively, with median shown as black lines. Significance of empP *<*0.05, after Bonferroni correction is shown with (*).

## 3 Discussion

In this study, we present a method for improved utilisation of the accumulating GWAS data, in order to facilitate mechanistic interpretation of identified disease-associated non-coding genomic variants. We developed a comparative genomic approach using conserved synteny of large genomic regions termed genomic regulatory blocks across entire vertebrate evolution, in turn providing an optimal list of distal regulatory variants for each gene. Thereby, we highly expand the number of SNP-gene associations with the ones acting through long-range genomic interactions, while at the same time avoiding the multiple comparison problem and thus inevitable p-value inflation. We achieve this balance of tested enhancer-gene pairs by focusing exclusively on the well documented evidence of genomic regions harbouring distal regulatory elements. A staggering number of 37,051 unique GWAS loci (158,611 locus-trait pairs) have novel target genes predicted by targPred, while the GWAS catalog continues to expand both in terms of the quantity of deposited studies, and their biological complexity, with no sign of reaching redundancy [57].

First, we provide an easily accessible web tool targPred (http://targpred.irb.hr) for querying publicly available GWAS variants that are located in the GRB regions of the genome. targPred enables researchers and clinicians to search by phenotypic trait, GWAS study or individual SNP and, when a SNP is located in a GRB region, the output is a genomic neighbourhood visualization and information on transcriptional activity of genes associated with the locus harbouring the SNP (Figure 3). As demonstrated in the *BDNF* example (Figure 7), this information can be used in interpretation of particular genomic loci associated with phenotypes, to narrow down associations to be tested in individual *in vitro* experiments or by further diagnostic tests. We would like to stress that these associations are not, *per se*, a proof of an SNP-gene association, rather they guide further tests in an economical manner.

By comparing targPred to similar publicly available tools, we demonstrated that different approaches indeed often result in different associations. We first measured how well target gene lists produced by the original GWAS associations and targPred compare against two sets of ground-truth gene-trait associations: a set of annotated coding variants linked to individual traits (Figure 5A) [54], and a set of experimentally-validated regulatory variants associated to genes and traits through extensive scientific literature search (Figure 5B) [47]. These two comparisons point to various valid S2G assignments in both GWAS and the GRB-based predictions, however, the fraction of overlapping assignments are much higher in the Alsheikh set. This trend implicates that the long-range variant-gene assignments that have previously been documented in the literature will often be correctly reported in the GWAS catalog, but in case where no literature on the non-coding variant exists, other sources of knowledge, i.e. the datasets of known coding variants like ClinVar are underutilised, albeit often correctly detected by targPred. Moreover, despite a rather small novelty contribution demonstrated on the Alsheikh set of regulatory variants, the prostate carcinoma locus association to the *SOX9* gene, missed by the GWAS shown above, further demonstrates that the validity of the novel S2G assignments has to be evaluated on the locus-by-locus level.

We further compared to the S2G predictions produced by targPred against a range of state-of-the-art prediction tools: ABC, GeneHancer and cS2G (Figure 6). While these comparisons can be performed mostly on the quantitative level of predictive scores, they show that, when target gene predictions exist for the same SNP-trait association, the high scores of these methods are significantly overrepresented in GRB target genes predicted to be significantly activated by the regulatory variant over long-range interaction with that gene. However, it is worth noting here that none of the three compared methods have the comprehensive range of enhancer-promoter pairs as targPred, which is based on patterns of extreme non-coding conservation over the evolution of vertebrate genomes.

The case of the *BDNF* locus presented in Figures 6-7 exemplifies the interpretation process using targPred and the biological utility of proposed alternative (and potentially corrected) associations to downstream experimental work. In this GRB, the *BDNF* gene is predicted as the target gene sensitive to regulatory inputs over long-range. targPred indeed identifies a significant increase in transcriptional levels of the *BDNF* especially under the influence of the Enh5. The regulatory variant in genomic locus overlapping Enh5 has been originally assigned to *LINC02758* gene [55]. The ABC method does not report any enhancers in this genomic locus, nor does cS2G (linking enhancers in the other locus in this GRB to *METLL15* and *KIF18A*). The GeneHancer tool covering the broadest set of enhancers including the ones created by the FANTOM5, associates Enh5 to *METLL15* and *KIF18A* genes, albeit with low-confidence scores.

Finally, a comprehensive analysis of GWAS non-coding variants across hundreds of traits demonstrated comparatively higher propensity for localisation in regulatory elements and consequently associated regulatory variants for some diseases and phenotypic traits. To no surprise, a whole range of psychiatric disorders are found enriched, in line with previous functional over-representations found among GRB target genes, i.e. the genes under extreme long-range regulation (Figure 4C). However, other trends emerged like cancer, immune disorders, asthma as well as metabolic disorders. The mounting evidence from previous proof-of-concept studies in obesity [32] and schizophrenia [33], individual examples shown in this study for prostate cancer and childhood obesity, propensity for LRR for 3 out of 8 asthma ontology terms (Figure 4A) and over-representation of GWAS loci under LRR across entire lineage of psychiatric disorders (Figure 4B) warrant implementation of the proposed targPred method as a necessary step when interpreting GWAS SNP-trait-gene associations.

## 4 Methods

### 4.1 Transcriptional level enhancer-promoter associations

Enhancer–genes associations were quantified using cap analysis of gene expression (CAGE) data from the FANTOM5 project [58]. We modified the empirical p-value (empP) method previously published in [33] to include both activation and repression. The FANTOM5 dataset of 1,046 biosamples (cell lines, primary cells, and tissues), excluding time-series samples, was subset to 309 samples by merging technical and donor replicates via median transcripts-per-million (TPM) values. Replicates were identified by removing “donorX” or “poolX” identifiers from sample names.

For each FANTOM5-detected enhancer, candidate promoter genes within *±*6.4 Mb (maximal genomic regulatory block size) were evaluated. Per enhancer-gene pair, biosamples were stratified by enhancer activity: “E+” (enhancer CAGE signal *>* 0) versus “E0” (enhancer CAGE signal = 0). The observed test statistic was the signed median difference,

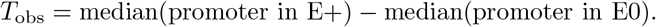

Null distributions were generated by randomly permuting biosample group labels (“E+”/”E0”) 10,000 times, recomputing *T*_perm_ each iteration. The two-tailed p-value tested bidirectional effects - activation and repression (|T_perm_| ≥ |*T*_obs_|):

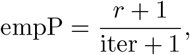

where *r* counts permutations at least as extreme as observed, and iter = 10,000. This pseudocount adjustment ensures empP *>* 0 and controls discreteness bias [59].

### 4.2 GWAS data processing

GWAS summary statistics files were downloaded from the GWAS catalog from the file repository with a data freeze on 17.02.2026. for all available studies present in the catalog at that time point [34]. All downstream processes were performed in R v4.3.3. Only studies done on European ancestry participants (defined in broad ancestral category) were used for downstream processing. SNPs that are involved in SNP-SNP interactions, multi-SNP haplotypes and SNP without a rsID were excluded. Studies that included multiple traits were separated so that each trait is considered separately. Furthermore, both reported and mapped genes were combined and hereafter termed GWAS genes. Calculation of linkage disequilibrium (LD) blocks was performed using the 1,000 Genomes phase 3 (GRCh37) dataset for the European population [60]. We selected canonical autosomal and X chromosomes and converted the data into bim/bed/fam format. The LD blocks were calculated for each SNP in GWAS data using PLINK v2.0 using its implementation of the exact solution to the cubic Hill equation [61]. The window for the LD block calculation was set to 500 kb and the *r*^2^ threshold was set to 0.6. For each GWAS SNP, a locus was spanning its’ corresponding LD block.

### 4.3 Colocalisation of GWAS loci with GRBs

GRB files and respective target genes were used from Tan, 2017 [62]. Briefly, human-dog pairwise comparison GRB files were used, based on CNEs with 98 sequence identity over 50 nucleotide window. Loci were defined as inside GRBs if there was any nucleotide overlap between two genomic ranges [33]. Target genes have properties which distinguish them apart from bystanders and other genes such as longer CpG islands, as well as more and wider spacing of alternative transcription start sites, and a specific combination of transcription factor binding sites in their promoter regions. The list of target genes was obtained from [62]. All other non-target genes located within a GRB were considered to be a bystander gene.

### 4.4 targPred web tool

To visualize the data on the web-based platform targPred, we generated two types of plots: genomic neighbourhood and two-sided distribution plots. The targPred online resource is based on the R Shiny v1.9.1 web application framework which enables an interactive user interface and also manages the server-side logic. The data tables are contained in a SQLite database, which is accessed using the RSQLite package v2.3.7 to dynamically query and retrieve relevant genetic information. The tables are rendered using the DT v0.23 package which allows fast pagination, searching, and sorting of the displayed data.

#### 4.4.1 Genomic neighbourhood

Genomic neighbourhood plots were created using R package Gviz v1.40.1. for each GWAS locus overlapping a GRB. It consists of nine automatically produced tracks, descriptions can be found in Table 1.

**Table 1.** Tracks in the genomic neighbourhood and their descriptions.

| Track | Description |
| --- | --- |
| Ideogram | Depicts the position of the neighbourhood on the chromosome. |
| Genome Axis | Marks the genomic ranges. |
| GRB | Depicts the genomic span of the overlapping GRB and names the target gene. |
| CNE density | Depicts density of conserved non-coding elements (CNEs) in the GRB. |
| Directionality index | The Homer human neural directionality index [63]. |
| Loci | Genomic span of the overlapping GWAS loci (i.e. LD blocks). |
| SNP | Positions of GWAS SNPs in their respective LD blocks. |
| eRNA | Positions of all eRNA enhancers identified using CAGE data that overlap the given locus. |
| Gene | Genes found in the GRB, where colouring scheme denotes their classification with respect to the GRB role. |

#### 4.4.2 Two-sided distribution plots

The two sided distribution plots were created using the R package beanplot version 1.3.1 and depict matrices of expression profiles (in log-transformed TPMs) across 309 biosamples, of genes in interaction with their respective enhancers. The gray part of the plots depicts the gene’s expression profiles in those biosamples where the enhancers are inactive (“E0”), while the pink part depicts the expression profiles for biosamples with actively expressed enhancers (“E+”). The horizontal lines denote the median values of the expression profile and the asterisk signs denote the level of significance for the association of the enhancer-gene pair. The significance level was controlled for multiple testing for all enhancer-gene pairs in a GRB using Bonferroni adjustment of empP.

### 4.5 Ground-truth SNP-gene associations

A set of credible trait-associated genes was curated based on coding variants deposited on ClinVar [54]. The VCF files were downloaded from the NCBI public repository site (data freeze 09.03.2026.) [64] and processed to exclude entries with unspecified diseases. Disease ontologies were used to cross-map coding variant disease annotation with EFO ontology which is used in GWAS annotations. A small number of direct mapping was possible and cross-mapping of synonyms was performed with MONDO ontology [65]. The final set consisted of trait-gene mappings for traits that were successfully mapped to GWAS traits. The second set of credible S2G was obtained from the original publication [47]. Only SNP-trait pairs which are found in GWAS and overlap GRBs were considered.

### 4.6 Enhancer-gene predictive methods

Genome wide ABC scores calculated across 131 biosamples were downloaded from the official website (https://mitra.stanford.edu/engreitz/oak/public/Nasser2021). The GeneHancer scores generated upon publication were obtained by request via email [28]. Both datasets were filtered for enhancers which overlap GRBs. Each enhancer-gene pair from the datasets was assigned to a category 0, 1 or 2 based on targPred. Category 0 are pairs which do not include a empP significant pair from our dataset nor the gene is a GRB target gene; category 1 are pairs which include either an empP significant pair from our dataset or a GRB target gene; category 2 are pairs which are both an empP significant pair from our dataset and a GRB target gene. Due to discontinuous values of GeneHancer scores (Figure 6C) only pairs with values > 250 were considered for testing. SNP-trait-gene triplet cS2G scores (published in [29]) for GWAS SNPs were downloaded from the repository (https://zenodo.org/records/7754032). We assigned a locus to each SNP which falls into a GRB, and followed the same protocol of assigning categories as before.

### 4.7 Traits under LRR

Identification of traits under LRR was performed by comparing loci-associated enhancer-gene pair which overlap GRBs versus those that fall outside of GRBs. First, for each trait, loci-associated enhancers were identified and categorized as inside or outside GRBs. Second, for each enhancer, a set of significant enhancer-gene pairs was identified (empP < 0.001). We have used raw p-values (uncorrected for multiple testing) while enforcing a more stringent threshold than elsewhere where correcting for multiple testing, since it is not possible to correct on the basis of a GRB region for those enhancer-gene pairs which are outside of GRBs. The median of genomic distance for each enhancer-gene pair was calculated and the difference in median distance between the two groups was tested using Wilcoxon rank sum test and corrected for false discovery rate using BH procedure. Additionally, traits were categorized into higher terms using EFO or MONDO ontologies.

The two EFO terms “overnutrition” (MONDO:0003916) and “diabetic eye disease” (EFO:0009486) have been placed in the aetiologic and body system subtrees and both lack systematization by developmental or physiological process, within the EFO ontology. We have therefore included these into the metabolic class of phenotypes in our analysis, but mark them in italic to differentiate from true child terms of “metabolic disease” (MONDO:0005066).

### 4.8 Locus enrichment in GRBs

Enrichment of trait-associated loci was performed by genome shuffling of GRB regions using bedtools v2.30. The shuffling was performed 10,000 time using GRBs and by specifying that the shuffled regions are not allowed to overlap each other and the original GRBs. To calculate the enrichment of loci in GRB, we selected GWAS studies with at least 30 associated SNPs. The number of overlaps within each of the 10,000 shuffled GRBs were counted to obtain a null distribution. The p-value was calculated empirically by counting how many times a value (i.e. counts) as extreme as the observed appeared in the 10,000 permutations.

### 4.9 Childhood obesity loci example

Childhood obesity region chromatin contact map was generated from Hi-C results on fully differentiated human fibroblast cells using 10,000 base-pair resolution [56]. Lift-over from hg38 to hg19 of the region was performed using R library rtracklayer v1.54. The visualization of the genomic region was generated using R library plotgardener v1.0 and two-sided distribution plots were obtained from the targPred data.

## Supplementary information

Supplemental table 1 - result of Wilcoxon test for number of enhancers for target and bystander genes per genomic distance bin.

Supplemental table 2 - result of locus permutation test for 4 psychiatric and 4 metabolic traits.

Supplemental table 3 - comparison between targPred and validated SNP-gene lists.

Supplemental table 4 - comparison between targPred and GeneHancer, ABC and cS2G.

Supplemental table 5 - scores and targPred categories for GeneHancer, ABC and cS2G.

Supplementary figures - Supplementary figures S1 and S2.

## Declarations

## Acknowledgements

Authors would like to thank dr. Tomislav Šmuc for insightful discussions during research inception and implementation, and guidance during this installation grant. This research was supported by the Croatian Science Foundation project “Exploring Interactions Between Regulatory Variants in Human Disease Context (IntRegVar)”, UIP-2020-02-1623.

## Funding

This study was conducted as a part of the “Exploring Interactions Between Regulatory Variants in Human Disease Context” project (UIP-2020-02-1623) funded by the Croatian Science Foundation. Katarina Mandić is funded by the Croatian Science Foundation project DOK-2021-02-7831.

## Conflict of interest/Competing interests

The authors declare no competing interests.

## Ethics approval and consent to participate

Not applicable.

## Consent for publication

Not applicable.

## Data availability

Not applicable.

## Materials availability

Not applicable

## Code availability

https://github.com/mlkr-rbi/targPred

## Author contribution

A.B. has designed the study and acquired funding for this work. B.L. has participated in conceptualising this research, and helped build the targPred method. K.M., D.H. and F.U. have built the web resource and performed all the analyses. A.B. and K.M. wrote the first version of the manuscript. All authors contributed to the writing of the final manuscript. All authors read and approved the final manuscript.

## Notes

### Competing Interest Statement

The authors have declared no competing interest.

