## Supplemental figures for "The role of long-range transcriptional regulation in interpretation of non-coding variants associated with human disease"

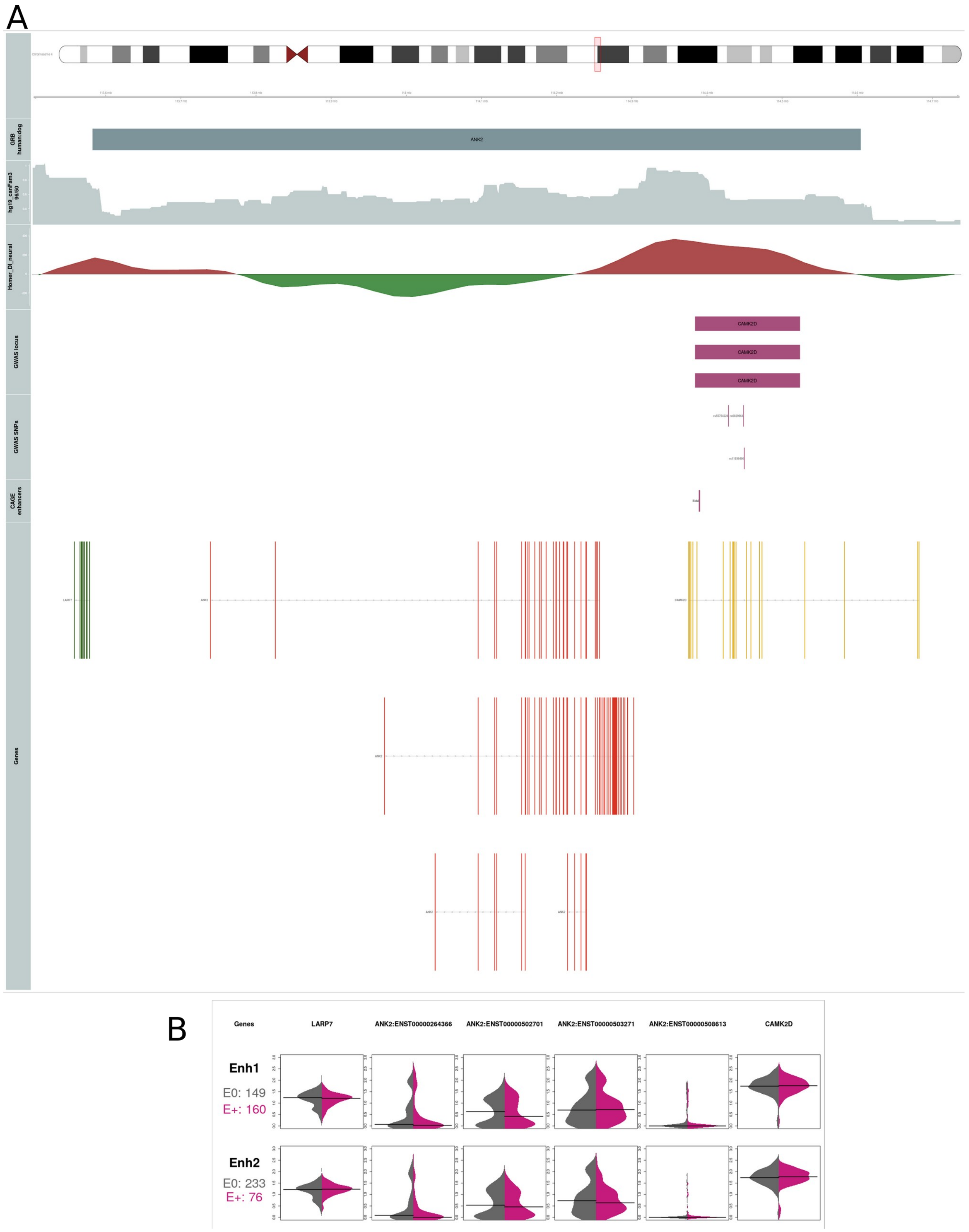


**Supplemental figure S1 A** Genomic neighbourhood of the GRB (chr4:113582644-114604487) with the target gene ANK2. Locus (chr4:114384328-114523729) contains two variants (rs55754224, rs6829664) associated with atrial fibrillation for which the GWAS associatedss CAMK2D gene. **B** Two-sided expression matrix consisting of enhancers overlaping the locus and genes found in the region.

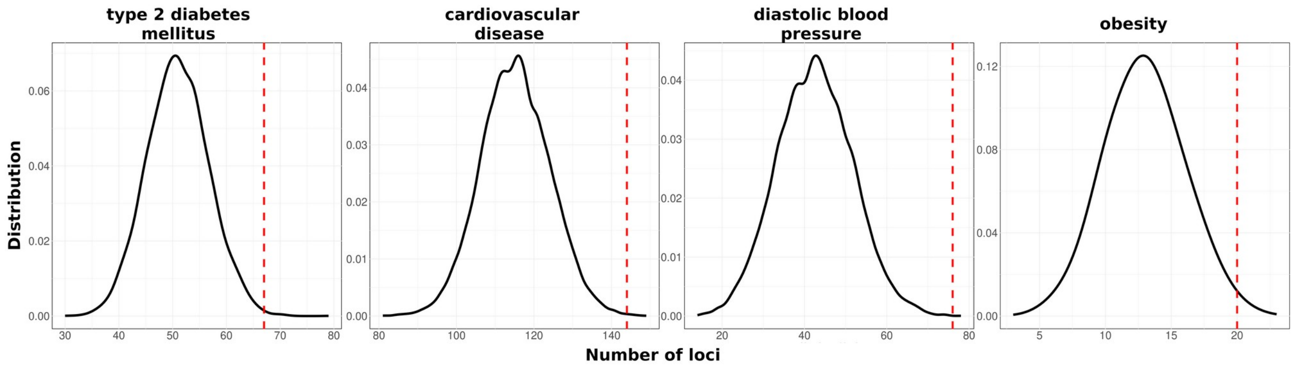


**Supplemental figure S2** Probability distribution of loci overlapping N = 10,000 permutated GRB regions was estimated for four traits associated with the metabolic syndrome. P-values were calculated empirically (Supplemental table 2).
